# Visualization of conformational changes and membrane remodeling leading to genome delivery by viral class-II fusion machinery

**DOI:** 10.1101/2022.03.12.484076

**Authors:** Vidya Mangala Prasad, Jelle S. Blijleven, Jolanda M. Smit, Kelly K. Lee

**Affiliations:** Department of Medicinal Chemistry, University of Washington, Seattle, WA 98195, USA; Zernike Institute for Advanced Materials, University of Groningen, Groningen, The Netherlands; Department of Medical Microbiology and Infection Prevention, University of Groningen, University Medical Center Groningen, Groningen, The Netherlands; Biological Physics, Structure and Design Graduate Program, University of Washington, Seattle, WA 98195, USA; Department of Microbiology, University of Washington, Seattle, WA 98195, USA

## Abstract

Chikungunya virus (CHIKV) is a human pathogen that delivers its genome to the host cell cytoplasm through endocytic low pH-activated membrane fusion mediated by class-II fusion proteins. Though structures of prefusion, icosahedral CHIKV are available, structural characterization of virion interaction with membranes has been limited. Here, we have used cryo-electron tomography to visualize CHIKV’s complete membrane fusion pathway, identifying key intermediary glycoprotein conformations coupled to membrane remodeling events. Using sub-tomogram averaging, we elucidate features of the low pH-exposed virion, nucleocapsid and full-length E1-glycoprotein’s post-fusion structure. Contrary to class-I fusion systems, CHIKV achieves membrane apposition by protrusion of extended E1-glycoprotein homotrimers into the target membrane. The fusion process also features a large hemifusion diaphragm that transitions to a wide pore for intact nucleocapsid delivery. Our analyses provide comprehensive ultrastructural insights into the class-II virus fusion system function and direct mechanistic characterization of the fundamental process of protein-mediated membrane fusion.

## Introduction

Chikungunya virus (CHIKV) is a mosquito-borne human pathogen that has caused major outbreaks in Europe, Asia and the Americas^1; 2^. It is a member of the alphavirus genus in the *Togaviridae* family^3^. Along with other members including Ross River virus, Semliki Forest virus, Sindbis virus and Venezuelan equine encephalitis virus, alphaviruses are responsible for severe emerging diseases in humans and animals^1; 4; 5^. CHIKV infections are characterized by high fever, fatigue, joint and muscle pains, with serious long-term effects including debilitating polyarthralgia^6; 7^. Despite its medical importance, no vaccines or antivirals against any alphavirus is currently available^8; 9^.

CHIKV, like all alphaviruses, is a membrane-enveloped, single-stranded, positive-sense RNA virus with an ∼11.8kb genome ^3; 10^. The mature CHIKV virion is composed of an icosahedral inner nucleocapsid containing 240 capsid monomers that enclose the viral genome^11^. The nucleocapsid is surrounded by a membrane bilayer. The external surface of the mature virus contains 240 copies of E1 and E2 membrane-anchored glycoprotein heterodimers, arranged as 80 trimeric spikes following icosahedral symmetry^3; 10; 11^ (Figure 1a-c). E2 is primarily responsible for cellular receptor attachment ^12; 13^ but also interacts non-covalently with the nucleocapsid to stabilize the virion structure ^14^. The E1 glycoprotein contains the hydrophobic fusion loop (FL) and mediates membrane fusion ^15; 16^. In the mature virion, E2 is positioned above E1, shielding the functionally critical FL from premature exposure^11; 17^ (Figure 1b,c).

**Figure 1.**
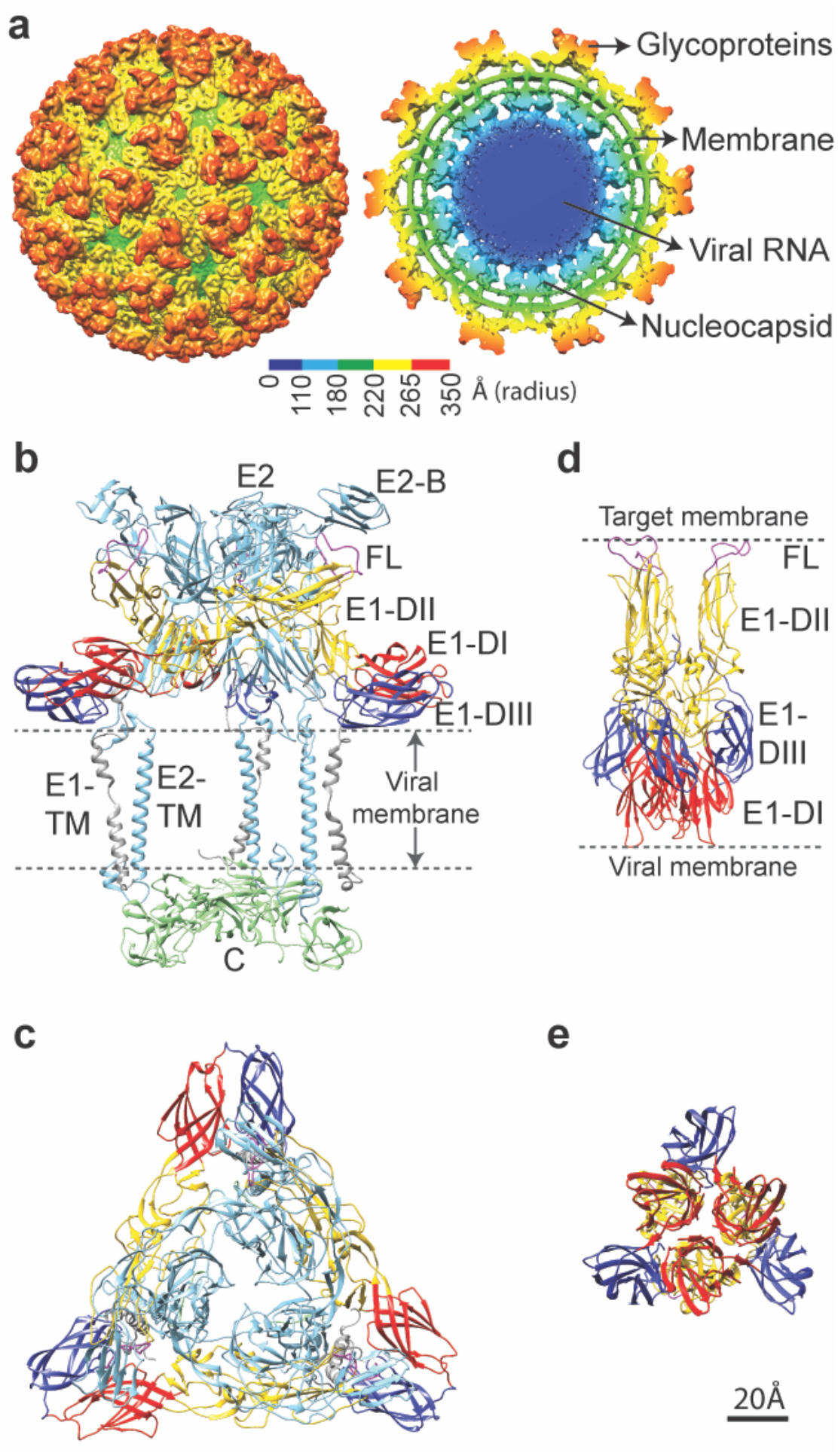
Structure of CHIKV particle. **a**. Surface view (left) and cross-sectional view (right) of UV-inactivated CHIKV strain S27. Cryo-EM density map is colored according to radius. **b**. Side view of ribbon structure of the trimeric surface glycoprotein heterodimers in contact with the inner capsid protein as observed in the wild-type virion (PDB ID: 3J2W). **c**. Top view of the trimeric E1-E2 heterodimers on wild-type CHIKV. **d and e**. Side and top view of the crystallographic structure of post-fusion E1 homotrimers (PDB ID: 1RER) respectively. In all panels, E1 is colored according to domains (domain I: red, domain II: yellow, domain III: blue, FL: magenta, E1-transmembrane domain: grey), E2 is in light blue and capsid protein in green.

CHIKV enters host cells primarily via clathrin-mediated endocytosis^18^ following attachment to a cellular receptor such as MxRA8^19^ or other attachment factors such as heparan sulfate or C-type lectins^20^. Upon cellular entry, the virus is engulfed into endosomes where the low pH environment resulting from endosomal maturation triggers conformational changes on the virus surface including the dissociation of the E1-E2 heterodimer^21^ and the formation of extended E1 homotrimers (HT) with its FLs inserted into the target membrane^22; 23^ (Figure 1d,e). The E1-HTs are then thought to drive membrane fusion by refolding/hairpin formation to bring the opposing membranes together^22^. Lipid mixing between the viral and endosomal membrane results in fusion pore formation that allows delivery of the viral nucleocapsid into the cytoplasm where it subsequently disassembles to release the viral RNA and establish infection^24; 25^.

The current model for how alphavirus membrane fusion takes place is primarily based on x-ray crystallographic structures of the pre-fusion^26^ and post-fusion conformations of recombinant E1 glycoprotein ectodomains^22^ along with related molecular studies on isolated glycoproteins^23; 27^. We lack direct structural data describing the sequence of protein conformational changes and nature of membrane remodeling that is necessary to derive a mechanistic understanding of the fusion process for alphaviruses and, more broadly, the type of fusion system (class-II) they represent^28^.

Here, we have used cryo-electron tomography (cryo-ET) in combination with sub-tomogram averaging to trap and observe the membrane fusion process in CHIKV under near-native conditions. Through stepwise analysis of CHIKV interactions with a target membrane at varying pH values and reaction timepoints, we can clearly demarcate intermediate stages in CHIKV membrane fusion. These data provide comprehensive insights into changes in virion structure, glycoprotein conformations, and changes in membrane organization along the fusion pathway. Our results also demonstrate that membrane fusion mediated by class-II fusion proteins in CHIKV proceeds by a markedly different pathway relative to class-I viral fusion systems such as influenza virus^29; 30; 31; 32^. Furthermore, our results highlight the power of cryo-ET for capturing 3-dimensional snapshots of reaction intermediates in a dynamic biological process from start to finish.

## Results

For our experiments, CHIKV (strain S27) particles were rendered replication incompetent by UV-light inactivation. The UV-treated virus drives membrane fusion in a similar fashion to untreated virus^33^. Single particle cryo-EM reconstruction of the UV-treated CHIKV was calculated to a resolution of 6.75 Å, which confirmed that the virion structure at neutral pH is identical to reported CHIKV structures^11; 19^ (Figure 1a, Supplementary Figure 1).

CHIKV particles were mixed with liposomes at varying pH conditions and incubated for a range of time points prior to plunge freezing in liquid ethane. Liposomes were prepared based on previous reports for optimal fusion in CHIKV^33^. At pH 6.5 and below, rapid aggregation of CHIKV particles was observed, hence, optimization of the ratio of liposomes to CHIKV was performed to reduce aggregation. The pH threshold for CHIKV S27 fusion is 6.2 with optimal fusion occurring in the pH range of 4.5-5.6^33^. Within this range, most particles carry out fusion within 10 seconds of exposure to low pH at 37 ºC^33^, exhibiting similar kinetics to other alphaviruses^34; 35^. To better sample and capture intermediate fusion stages within the constraints of cryo-EM grid preparation conditions, membrane fusion experiments were performed at room temperature, which slows the fusion reaction.

At pH values above 6.0, CHIKV particles associated with liposomes via discrete densities bridging the virus-liposome interface (Supplementary Figure 2). However, interactions beyond the initial virus-liposome association were rarely observed even at longer incubation periods of ∼30 minutes. Indeed, even in fluorescence studies, the extent of fusion events observed at pH 6.0 and above was negligible^33^. We observed a clear progression to completion of fusion only at pH values below 6.0. At pH ≤5.0, even at room temperature, most virions in the population completed the membrane fusion process within 15 seconds. Thus, for better sampling of fusion events, pH values in the intermediate range including 5.9, 5.6 and 5.1 were examined. Across these pH values, the observed intermediates are similar, except that at lower pH, a more rapid progression through steps leading to complete fusion was observed. From analysis of more than six hundred CHIKV-liposome complexes in our tomograms, CHIKV-mediated membrane fusion stages could be categorized into nine distinct steps, which are discussed in detail below.

### Stage I - Membrane recruitment

Initial membrane recruitment can be observed between 30 seconds-1 minute at pH 6.1 and 5.9, and within 30 seconds at pH 5.6. Minute, localized attachments are observed, sparsely bridging the CHIKV glycoprotein exterior to the liposomes, with the glycoprotein shell appearing relatively intact (Figure 2a-c, Supplementary Video 1). From analysis of more than one hundred such interactions, the lengths of the delicate attachments extending from the viral glycoprotein surface to the liposome surface were observed to be ∼32-45 Å. At neutral pH on the virus surface, the E2 B domain protects the E1-FL from solvent exposure^11^ (Figure 1b), but under low pH conditions, the B domain has been reported to exhibit increased flexibility resulting in E1-FL exposure and potential for membrane binding^17; 36^. The fine attachments seen in the tomograms (Figure 2a-c) thus likely reflect a state in which the tip of individual E1s containing the FL have inserted into membrane, but without global disruption of the trimeric E1-E2 arrangement on the virus surface (Figure 2d).

**Figure 2.**
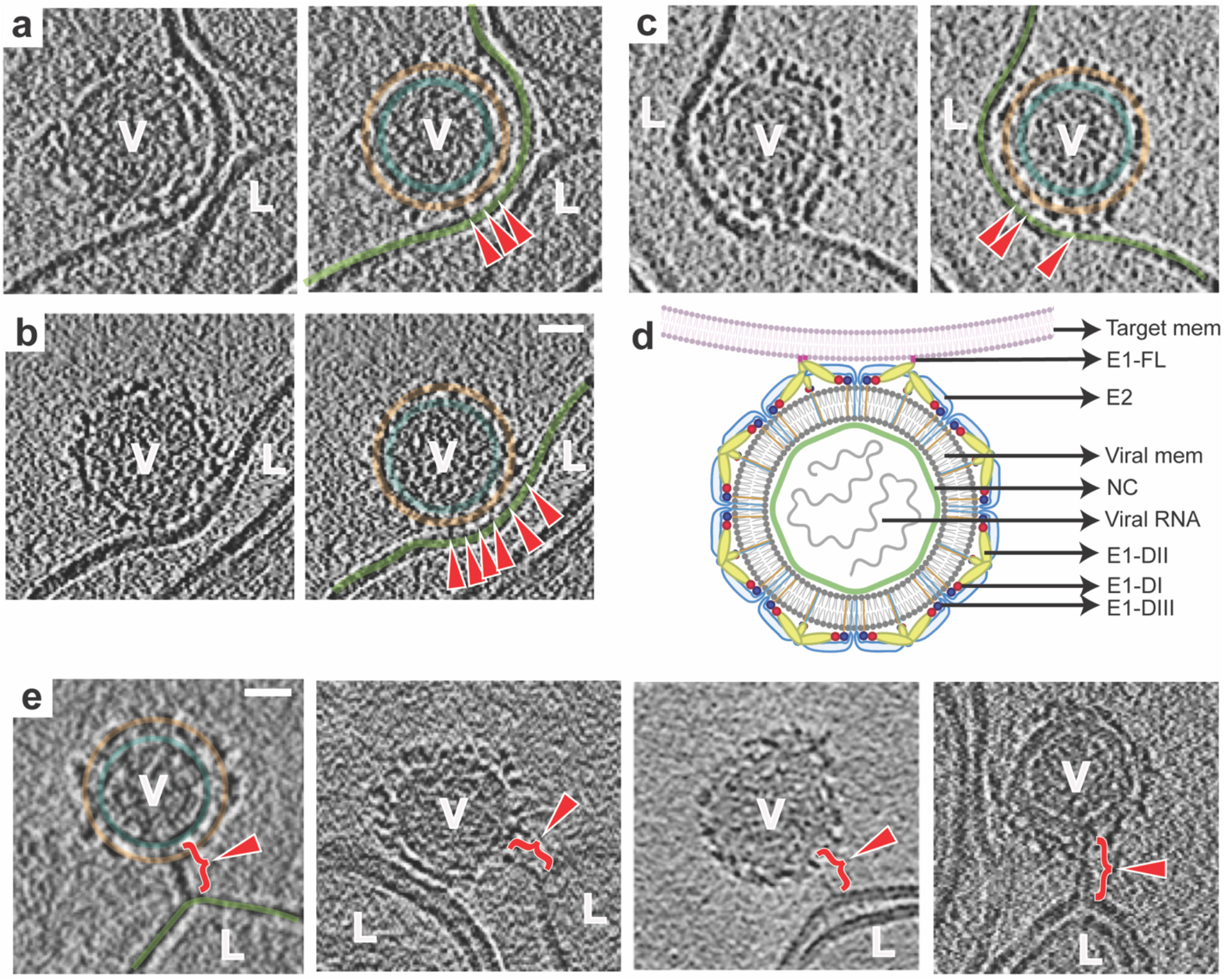
CHIKV membrane recruitment – stage I. **a-c**. Left panel: Tomogram slice showing CHIKV (V) interacting with liposome (L) via thin, delicate attachments. Right: Same tomogram slice as in the left panel but annotated to show the different protein and membrane layers: CHIKV glycoprotein layer in orange, CHIKV membrane in teal and liposome membrane in green. The attachments between CHIKV and liposome membrane are denoted by red arrowheads. **d**. Cartoon representation of this fusion stage. **e**. Tomogram slices showing examples of singular, long glycoprotein connection (red bracket with arrowhead) to liposome membrane. Leftmost panel alone is annotated similar to previous panels for reference. Scale bar is 200 Å in all panels. Black is high density in all panels.

While the CHIKV surface appeared globally intact, sub-tomogram averaging of low pH (<6.0) exposed CHIKV particles that were isolated or weakly attached to target membranes, showed that the virions have deviated from their global icosahedrally symmetrical organization (Supplementary Figure 3D). Due to the low number of particles available, it was not possible to determine whether the virions retained local symmetry features. Comparison of 2D radial density plots of neutral pH CHIKV with the sub-tomogram averaged low pH CHIKV structure showed that the outer glycoprotein shell in low pH CHIKV had expanded by ∼20 Å relative to neutral pH CHIKV (Supplementary Figure 3).

### Stage II - Membrane attachment

The next stage of glycoprotein engagement with the target membrane is accompanied by a transition of E1 from its orientation parallel to the virus surface to a more perpendicular orientation with respect to the surface. This stage occurred by 1 minute at pH 5.9, by 30 seconds at pH 5.6 and almost instantaneously at pH 5.1.

In ∼2% of examples of CHIKV at early time points, singular, hyper-extended glycoprotein density was seen interacting with the target liposome. The connecting density in these cases were ∼170 Å - 250 Å as measured from the viral membrane surface to the liposomal membrane (Figure 2e). In contrast, crystal structures of the E1 ectodomain in its pre-fusion and post-fusion conformations have a length of only ∼125 Å^26^ and ∼100 Å^22^ respectively (Figure 1b,d). Thus, these extended connections are only feasible with major changes in the E1 structure involving hyperextension and repositioning of component domains. This also suggests that the energy needed to detach E1 from the target membrane is larger than that required to partially unravel the E1 subunit.

More commonly at this stage, extensions of clustered glycoprotein density and formation of multiple robust attachments were observed between the glycoproteins and the liposomes at the interaction interface (Figure 3a-c, Supplementary Video 2). For virion facets that were not interacting with membranes, heterogeneity in glycoprotein organization was evident on the particle exterior. From analysis of 221 interaction sites, consisting of multiple glycoprotein attachments to liposomes, the length of these connecting densities ranged between 90 Å to 165 Å on central tomogram slices (perpendicular to electron beam direction), as measured from the viral membrane surface to the target membrane. Corroborating our observations, long, bridge-like densities, attributed to the E1 protein, have been imaged previously in an early-stage fusion intermediate of Sindbis virus in contact with liposomes^37^. Furthermore, in our tomograms, residual protein density was seen close to the viral membrane at the virus-liposome interface (Figure 3a-c) suggesting that the E2 proteins may still be present at the particle-liposome interface, similar to that seen with Sindbis virus^37^. The observed multiple attachments between E1 and liposome membrane appear to set the stage for further steps that involve concerted action of multiple E1 proteins.

**Figure 3.**
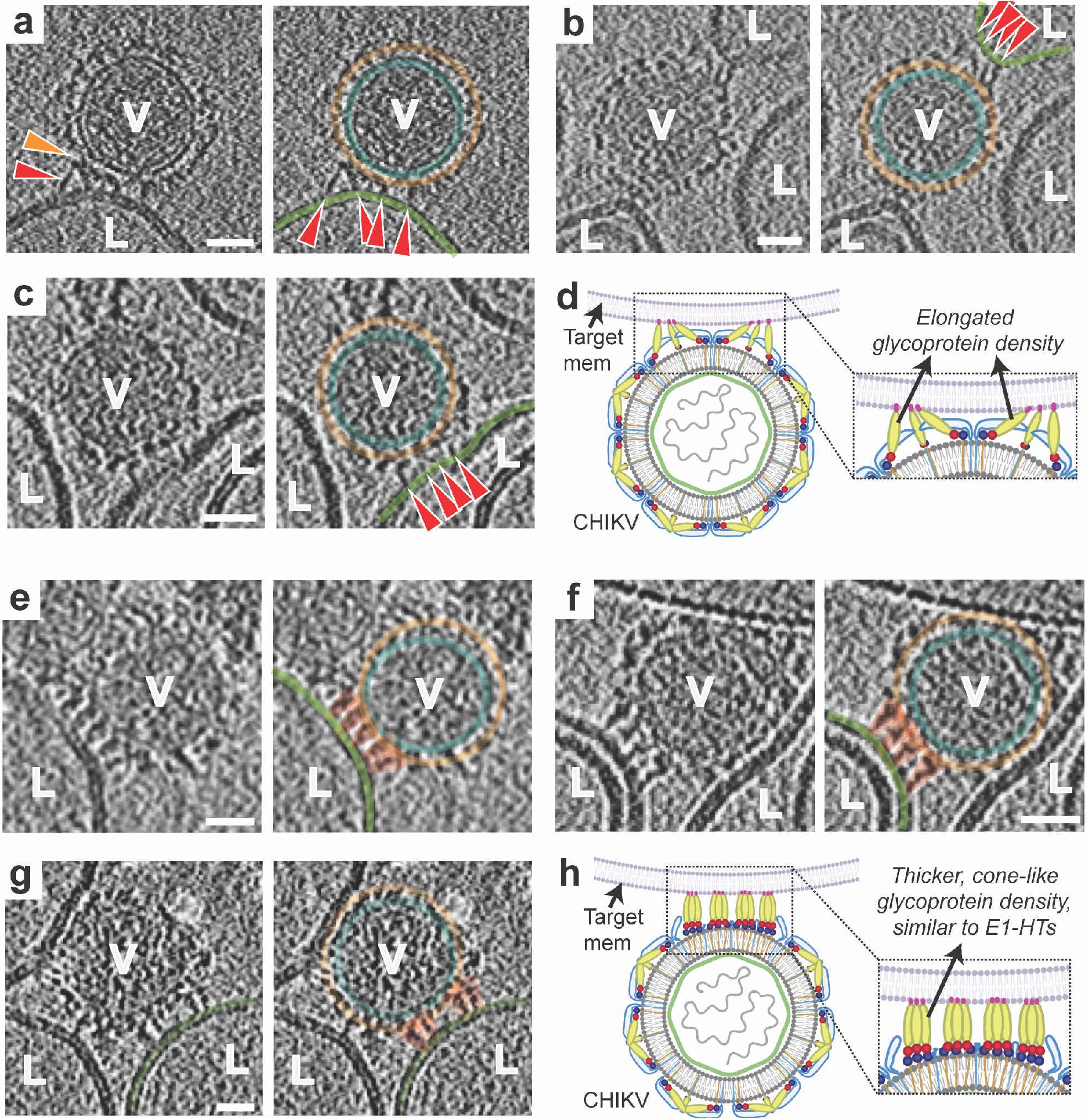
Glycoprotein membrane attachment and E1 homotrimer formation (stages II and III). **a-c**. Stage II. Left panels: Tomogram slice showing long bridge-like attachments between CHIKV (V) and liposomes (L). Red arrowhead indicates extended glycoprotein density and orange arrowhead denotes remaining glycoprotein density close to the viral membrane surface. Right: Same tomogram slice as in the left panels but annotated to show the different protein and membrane layers. CHIKV glycoprotein layer in orange, CHIKV membrane in teal and liposome membrane in green. Glycoprotein attachments between CHIKV and liposome membrane are denoted by red arrowheads. **d**. Cartoon representation of this fusion stage with zoomed inset showing the virus-target membrane interaction interface. **e-g**. Stage III. Similar representation as in a-c with left panels showing the raw tomogram slices and right panels showing the same slices with annotation. Cone-like glycoprotein densities that resemble E1-HTs are colored in orange. **h**. Cartoon representation of the E1-homotrimer formation stage with zoomed inset highlighting the region of interest. Scale bar is 200 Å and black is high density in all panels.

### Stage III – E1 homotrimer (HT) formation

Once multiple attachments between E1 and target membrane have formed, the E1 glycoproteins at the virus-target membrane interface transition to form thick, cone-like densities that are perpendicular to the viral and target membrane planes (Figure 3e-g, I, Supplementary Video 3). These features (Figure 3e-g) are similar in shape to the known crystal structures of post-fusion E1 trimers (Figure 1d,e) suggesting that the E1 proteins have oligomerized at this stage to a form of E1 homotrimers (HT). Four to five E1-HTs can be identified clustered at a given virus-liposome interface (Figure 3e-g). The E2 proteins appear to have been displaced from the virus-liposome interface to allow E1 trimerization. E1-HTs were observed by 2 minutes at pH 5.9 as well as pH 5.6 and by 30 seconds-1 minute at pH 5.1.

At the resolution of our cryo-ET data, it is not possible to directly discern whether swapping of domains I and III of E1, as seen in the crystal structures of post-fusion E1 ectodomain trimers ^22^, has occurred. The lengths of E1-HTs in our tomograms are ∼130-150 Å whereas the length of the post-fusion E1-trimers from crystal structures measures ∼100 Å (Figure 1d). The E1-domain III is ∼30Å in dimension. The cryo-ET data, thus, indicate that the domain III of E1 has likely not folded back to produce the post-fusion conformation at this stage. Our inference regarding this extended E1-HT state is supported by the identification of a pre-fusion intermediate form of E1-HT in previous molecular studies with Semliki Forest virus and Sindbis viruses^27; 35^.

### Stage IV - E1-HT membrane insertion

Once extended E1-HT formation occurs, the trimer appears to drive through the target membrane, causing depressions and possibly small punctures to the target membrane integrity (Figure 4a-e, Supplementary Video 3,4). This stage of E1-HT membrane insertion can be observed by 2-5 minutes at pH 5.6 and by 30 seconds-1 minute at pH 5.1. Supporting our observations, insertion of purified low pH-induced E1 homotrimers (full-length and ectodomain) into liposomal membranes has been previously reported ^22; 38^. Exact measurements of glycoprotein length in this stage were challenging owing to interference from surrounding membrane density. However, in cases where measurements could be made, such as in examples shown in Figure 4b-d, the E1-HT length varied from 110-150Å suggesting that complete folding-back of E1-domain III had still not occurred. In our cryo-ET data, we also observe examples of neighboring glycoprotein densities attaching to the target membrane (Figure 4b-d). With increasing numbers of glycoprotein attachments to the target membrane, the membrane is pulled towards the virion and can be observed to follow the contours of the virion exterior (Figure 4b-d). Projection of E1-HT into the target membrane at these closely packed interfaces appears to be responsible for bringing the target membrane close to the viral membrane.

**Figure 4.**
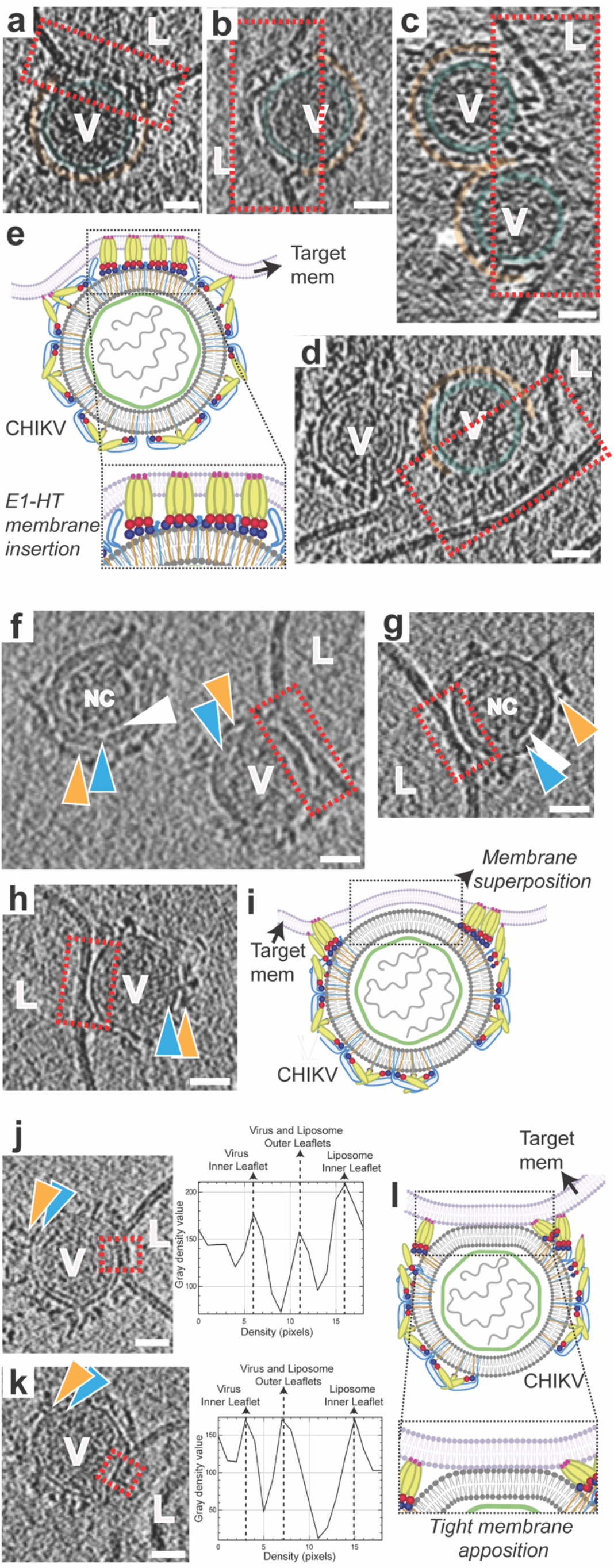
E1-HT membrane insertion and opposing membrane superposition (stages IV-VI). **A-d**. Annotated tomogram slices showing insertion of protein densities into the liposome (L) causing disruption of membrane density at the interface. CHIKV (V) are annotated similar to previous figures with red dotted rectangles enclosing interaction areas of interest. **e**. Cartoon representation of fusion stage IV – E1-HT membrane insertion. **f-h**. Tomogram slices showing superposition of the viral and liposome membranes. Interaction interfaces are enclosed in red rectangles. Variation and fluidity in the glycoprotein layer (orange triangles) can be seen in panels f,g. White triangles represent the gap observed between the NC and inner surface of the viral membrane (blue triangles). **i**. Cartoon representation of fusion stage V – membrane superposition, with dotted rectangle outlining the interface. **j-k**. Tomogram slices showing tightly docked membrane interfaces with the proximal leaflets too close to separate at the current tomogram resolution. Corresponding electron density plots along a line traversing the tight-membrane interface in the boxed region (red) of the tomogram slices are also shown. **l**. Cartoon representation of the tightly docked membrane interface (stage VI) with zoomed inset showing region of interest. Scale bar is 200 Å and black is high density in all panels.

### Stage V and VI - Opposing membrane superposition

In similar timepoints as E1-HT membrane insertion, opposing membrane superposition was also observed. As the membrane-inserted conformation of E1-HTs are not a favorable condition for the predominantly surface exposed E1 proteins, we deduce that the E1-HTs likely are driven away from the virus-liposome interface, resulting in their exclusion from the contact zone, which instead contains the two membranes in close proximity to each other with only an ∼1 nm gap between the proximal leaflets (Figure 4f-i, Supplementary Video 5).

At these intermediate stages, starting from the stage of extended E1-HT formation, we observe glycoproteins being displaced laterally on the virion surface (Figure 4f,g). This indicates that the cytoplasmic tails of the E2 glycoproteins have been uncoupled from the internal nucleocapsid, affording them mobility that is restricted in prefusion CHIKV and early fusion stages. In a few cases, a larger gap between the nucleocapsid and viral membrane is seen (Figure 4f,g) with the nucleocapsid no longer juxtaposed against the inner side of the viral membrane.

At this stage, we concurrently also observed cases where the viral and target membrane bilayers were tightly docked together, such that the individual proximal leaflets were indistinguishable at the resolution of our tomograms, resulting in a distinct 3-layer membrane interface (Figure 4j-l, Supplementary Video 6). Such a configuration requires dehydration of the proximal leaflets to permit close approach and meshing of the polar headgroups^39^. Most likely the energy released from the surrounding glycoprotein activity is transduced into formation of this lipidic organization^40^. Similar tightly-docked membrane-membrane contacts have been reported previously as an intermediate during membrane fusion by influenza virus^30^ and intracellular SNARE proteins^39^.

### Stage VII - Hemifusion

Following the formation of tightly juxtaposed membrane interfaces, we next observe clear examples of merged outer leaflets of the viral and liposomal membranes, corresponding to hemifused membranes (Figure 5a,b, Supplementary Video 7). This stage is observed at 5 minutes at pH 5.6 and by 3 minutes at pH 5.1. It is possible that target-membrane insertion of E1-HTs followed by movement of E1-HTs away from the interface causes enough perturbation or strains in the membrane to encourage lipid mixing and merging of the proximal leaflets. At the junction between the viral and target membranes, we observed clear examples of E1-homotrimers that measure ∼100 Å (Fig. 5a), consistent with the size and shape of post-fusion E1 trimers^22^. This indicates that by the hemifusion stage, the extended E1-HTs have transitioned completely, with domain III folded back, to form post-fusion E1 trimers, driving tight membrane apposition and hemifusion.

**Figure 5.**
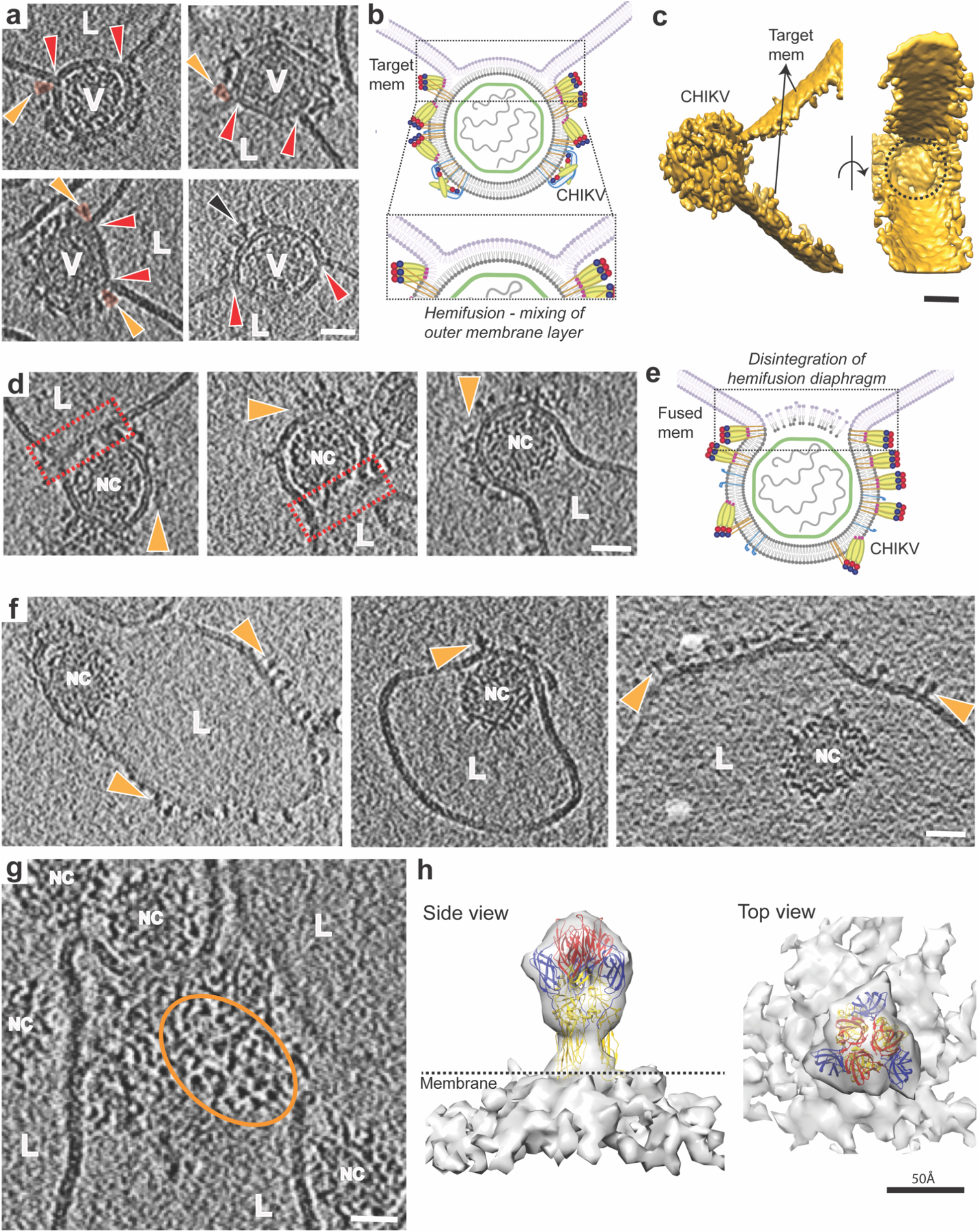
Membrane fusion stages VII-IX – hemifusion to nucleocapsid release. **a**. Tomogram slices showing clear examples of hemifusion or mixing of membrane leaflets between CHIKV (V) and liposomes (L). Red arrows indicate three-way junctions where the two membranes intersect. Orange arrows indicate glycoprotein density (also colored in orange) at the hemifusion junction that correspond to post-fusion E1 homotrimers. Black arrow shows presence of E1 homotrimers on virion membrane, suggesting that E1-FL can bind to viral membrane itself on availability. **b**. Cartoon representation of hemifusion with zoomed inset showing the region of mixing between the two outer leaflets. **c**. Surface 3D rendering of hemifused virion shown in bottom left of panel a. Side view (left) and 90° rotated view (right) is shown. The circular surface of hemifusion diaphragm can be seen (black dotted circle). **d**. Disintegration of the mixed central layer in the hemifused state leads to formation of a fusion pore. Fusion pore interface is shown as red rectangles and glycoproteins indicated as orange triangles. **e**. Cartoon representation of panel d. **f**. Subsequent release of the CHIKV nucleocapsid (NC) into the liposome lumen after fusion of the viral and liposome membranes. Floating glycoprotein densities on liposome surface are indicated in orange. **g**. Top view of a fused CHIKV showing triangle-shaped glycoprotein densities (orange oval) on the liposome surface. **h**. Sub-tomogram average of the glycoprotein densities seen in panels f and g, with the post-fusion E1-homotrimer crystal structure (PDB ID: 1RER) fitted into the density. Scale bar is 200 Å and black is high density in panels a-g.

Remarkably, the hemifusion diaphragm in CHIKV membrane fusion is quite large. In cases where this interface was resolved clearly in all directions, the diaphragm appeared almost circular with an average diameter of 350Å, which is nearly half the CHIKV diameter (Figure 5c). In general, hemifusion diaphragms ranged from half to full diameter of CHIKV, making them comparable in size to the nucleocapsid that needs to be delivered once the fusion pore forms (Figure 5a).

### Stage VIII – Fusion pore formation

Hemifusion in CHIKV progresses with disintegration of the hemifusion diaphragm, leading to formation of a fusion pore (Figure 5d,e; Supplementary Video 8). This stage is observed by 5 minutes at pH 5.6 and 3 minutes at pH 5.1. In agreement with the large hemifusion diaphragms, the fusion pore in CHIKV also exhibit widths >75% of the virion diameter (Figure 5d).

### Stage IX – Release of intact nucleocapsid

The last step of membrane fusion is the release of the CHIKV nucleocapsid (NC) into the liposome lumen. We observed more than 150 instances of NC released into the liposomal lumen and all of them appeared intact (Figure 5f,g; Supplementary Video 9). Nevertheless, the released NCs had lost icosahedral symmetry, as confirmed by our sub-tomogram averaging attempts of intact NCs. The presence of intact cores after membrane fusion confirm that further interaction with cellular host factors, such as the large ribosomal subunit ^24^, is required for nucleocapsid disassembly and release of the viral genome. The loss of icosahedral symmetry in the intact NCs further substantiates conformational changes in its structure as has been proposed previously to be necessary for exposing interaction sites that facilitate NC disassembly ^25^.

In the timepoints where membrane fusion has been completed and the intact NCs have been released into the liposome lumen, distinct protein densities decorate the exterior of fused liposomes (Figure 5f, Supplementary Video 9). These protein subunits originate at the virus-liposome fusion interface and are distributed across the entire fused liposome (Figure 5f). From top-down views of fused liposome surfaces in our tomograms, the protein subunits appear trimeric (Figure 5g). Sub-volumes of these protein subunits were extracted from the cryo-electron tomograms and subjected to sub-tomogram averaging. The resolution of the averaged structure is 27.2 Å at 0.5 FSC (Fourier Shell Correlation) cutoff (Supplementary Figure 4). The crystal structure of the post-fusion E1 homotrimer from Semliki Forest Virus (SFV) (PDB ID:1RER) fits in a unique orientation into the density map (Figure 5h), confirming that these protein subunits are indeed the post-fusion E1-trimers. Fitting the E1 ectodomain crystal structure into the density map shows that insertion of the E1 trimer into the outer membrane leaflet is only mediated by its FL without embedding additional regions of the ectodomain (Figure 5h). Moreover, no extra density is left to accommodate the E2 protein. These observations indicate that during and after membrane fusion, the E2 proteins do not form any oligomeric conformations and likely remain as individual protein subunits diffused across the membrane surface^21^.

### Other effects of low pH on CHIKV

At longer timepoints of pH 5.6 and 5.1, CHIKV can be often observed to undergo membrane fusion steps as a cluster of aggregated virions (Supplementary Figure 5a). Virions fused with each other suggesting that membrane attachment of E1 is non-specific (Supplementary Figure 5b). Furthermore, instances where one virion facilitated attachment and membrane fusion of an adjacent virion were also observed (Supplementary Figure 5a). Instances where CHIKV particles released NCs into the solution without any interaction with liposomes were also seen (Supplementary Figure 5b), suggesting that the CHIKV virion becomes increasingly unstable with decreasing pH.

## Discussion

Protein-mediated membrane fusion is a critical step in enveloped virus infection and a fundamental process that underpins many cellular functions. For viruses that employ class-II fusion proteins, such as flavi- and alphaviruses, virion architectures and structures of the pre- and post-fusion glycoprotein ectodomains are well established^28; 41^. However, as with most protein-mediated fusion systems, it has been challenging to obtain detailed structural information that describes the sequence of events that occur during membrane fusion in the context of the functional virion. Here, using cryo-ET, we have imaged the steps that an alphavirus must traverse during cellular entry under near physiological conditions. This approach has enabled us to identify multiple stages in the fusion process that were previously uncharacterized. By tracking the relative frequency of observed states at different time points (Figure 6a), the sequence of events leading to fusion and final release of the nucleocapsid was inferred (Figure 6b).

**Figure 6.**
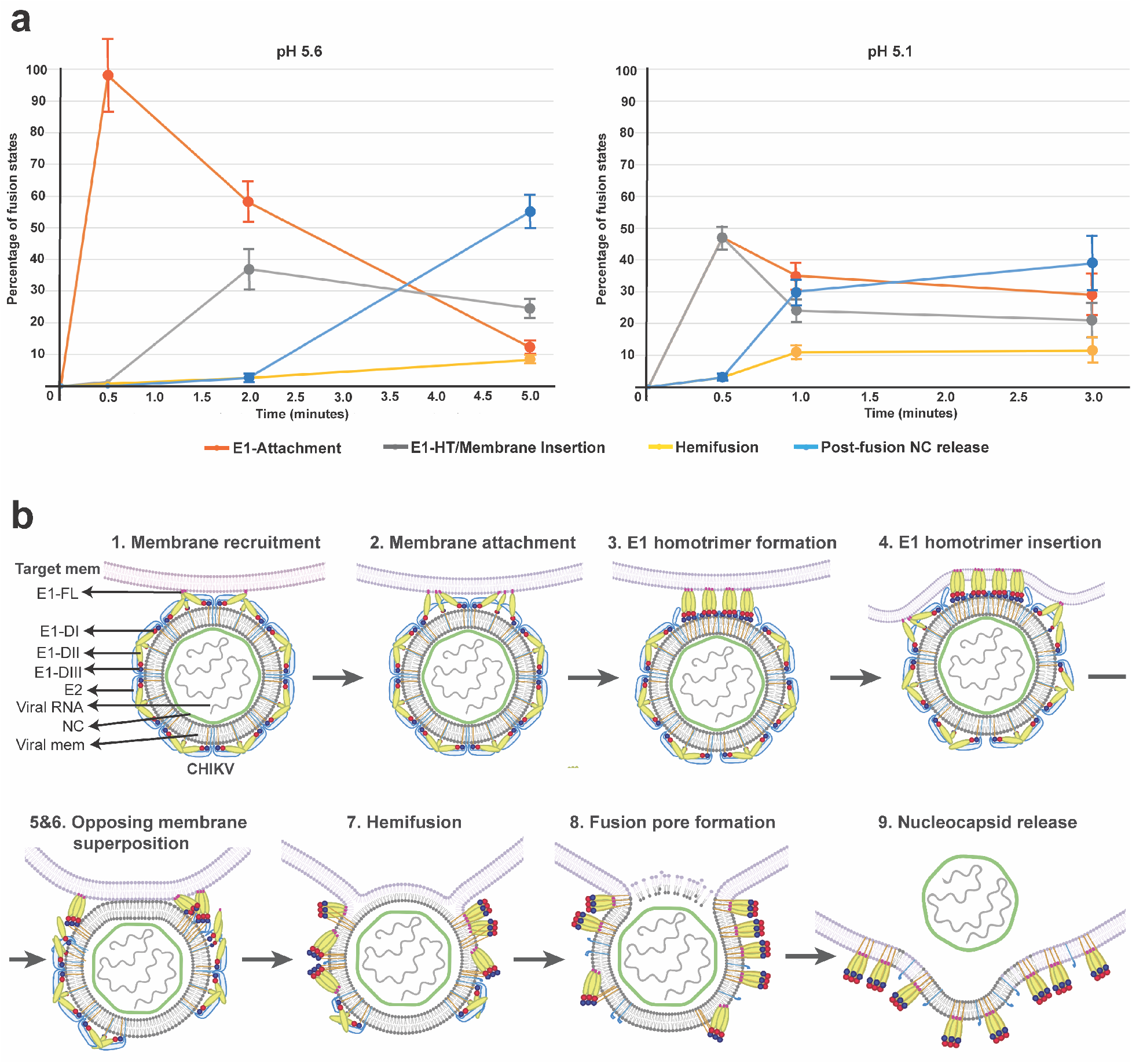
CHIKV membrane fusion pathway. **a**. Population of fusion-states during CHIKV membrane fusion. For pH 5.6 (left) and pH 5.1 (right), percentage of each fusion-state was calculated as a function of the total number of CHIKV-liposome contacts (n) observed at that given pH and timepoint. For pH 5.6, n=281 and for pH 5.1, n=263. E1 attachment (red plot line) includes stages I (membrane recruitment) and II (membrane attachment); E1-HT/Membrane Insertion (grey plot line) includes stage III-VI: E1-HT formation, E1-HT membrane insertion, membrane superposition and tight apposition; Hemifusion (yellow plot line) includes stages VII (Hemifusion) and VIII (fusion pore formation); Post-fusion NC release (blue plot line) includes stage IX (nucleocapsids released into liposomal lumen). Individual population counts for each state are given in Supplemental Table 1. Error bars have been calculated as square root of the number of complexes observed for each fusion state at the given pH and timepoint, similar to previous report^30^. **b**. Cartoon schematic of CHIKV membrane fusion stages.

Whereas class-I viral fusion proteins (such as influenza HA) adopt a trimeric prefusion conformation, class-II proteins in alpha- and flaviviruses are arrayed as symmetrically organized heterodimers and dimers on the prefusion virion^28; 40; 41^. Despite extensive quaternary interactions between E1 and E2 across the icosahedrally organized virion surface, the CHIKV E1 glycoproteins appear to be individually activated under the effect of low pH and membrane availability. This is similar to the case in influenza virus in which individual HA activate independently and adopt dynamic intermediate conformations^42^.

In our cryo-ET data, E1 monomers and homotrimers could be discerned in complete virions, placing class-II fusion protein intermediate structures that had been characterized as soluble, isolated components^22; 27; 38^ into the context of active membrane fusion reactions involving intact particles and target membranes. Our cryo-ET analysis further corroborates the role of an elongated form of E1-HT where its domain III has yet to fold back against the homotrimer core^27^. This elongated E1-HT state is somewhat analogous to the extended prehairpin intermediate conformations proposed for class-I fusion proteins^42; 43^. However, there is no evidence to date that extended pre-hairpin class-I trimers project into the target membrane as we observed with the CHIKV E1 proteins.

Folding back of E1 domain III leads to colocalization of the E1 membrane anchor and its FL on the same end of the protein leading to juxtaposition of viral and target membranes. Supporting this notion, in our data, we first observe presence of a shortened E1-homotrimer confirmation at the hemifusion stage, suggestive of E1-domain III fold-back and formation of the post-fusion E1 conformation. In previous cryo-ET studies of class-I fusion proteins, the extended intermediate of the glycoprotein trimer bends upon itself to bring the target membrane close to the viral membrane, leading to localized dimpling of the target membrane as it is drawn towards the viral membrane^28; 29^. That no such dimples were observed at any point in CHIKV class-II fusion pathway indicates that there are more pathways to effect protein-mediated membrane fusion than previously appreciated. In contrast, the extended, tightly docked membrane interfaces as seen in CHIKV (Figure 4j,k) have been observed during influenza virus^29^ as well as SNARE-mediated^38^ membrane fusion reactions. These observations underscore the generality of this membrane reorganization stage, indicating its role as an obligate intermediate state and highlighting its significance in protein-mediated membrane fusion reactions.

Once membrane apposition occurs, the CHIKV nucleocapsid detaches from the internal side of the viral membrane. This is likely necessary to provide fluidity to the viral membrane. Similar disintegration of matrix layers in influenza virus membrane fusion precede fusion pore formation^30^. In our study, we also observed that under optimal fusion pH conditions, the CHIKV virions, including its NC, lose their icosahedral nature. Furthermore, we observed slight expansion of the glycoprotein shell in the low pH-exposed CHIKV, consistent with reports for Semliki-forest virus at mildly acidic pH^44^. It is possible that acid-induced conformational changes in the surface glycoproteins are transmitted through the viral membrane to weaken the E2 glycoprotein’s cytoplasmic tail interaction with the NC^45^. Loss of interaction between the external glycoproteins and NC allows the glycoproteins to diffuse freely on the viral membrane, which enables direct interaction between the opposing membranes during the fusion process as is seen in our study (Figure 4). Changes in interaction between the E2’s cytoplasmic tail and NC has been implicated in causing structural changes in the NC^46^. Alternatively, ion leakage across the viral membrane either via the 6K protein^47^ or the E1 protein^48^ may permit acidification of the virus interior. It is possible that both the conformational changes in the glycoproteins and acidification of the virus interior mutually influence the NC structure. The present data, however, does not allow us to distinguish between the two possible mechanisms. Nevertheless, it is clear that the pH-dependent surface glycoprotein-NC protein interaction plays a key regulatory role in the alphavirus fusion system, much like the HA-matrix protein interactions observed in influenza virus^30^. This suggests that coordinated changes between the primary fusion protein and other structural proteins in enveloped viruses are a common phenomenon that likely help govern the sequence of membrane fusion events.

One key observation regarding lipidic intermediates relates to penultimate stages of membrane remodeling leading up to fusion pore opening. Notably, we observed several examples of hemifused diaphragms in our data. As the time-dependent evolution of intermediate populations shows (Figure 6a), a low fraction of hemifused complexes remain constant over time, even as examples of post-fusion complexes increase substantially. These observations suggest that hemifusion might represent a rate-limiting step in the membrane fusion process, as hypothesized via fluorescence studies of CHIKV membrane fusion^33^. In Ca2+ triggered membrane fusion reactions such as the SNARE-mediated systems, the hemifused configuration has been shown to embody a kinetically trapped state with productive fusion occurring instead through hemifusion-free point contacts ^49^. In our cryo-ET data, we do not observe any examples that suggest an alternate hemifusion-free pathway, though it is possible that such alternate pathways occur too rapidly to be detected in our cryo-ET conditions.

The present study provides the most detailed characterization of a class-II protein-mediated membrane fusion process by resolving protein intermediates and non-canonical membrane configurations associated with protein remodeling. These results chart the molecular processes that alphaviruses and other class-II fusion virus systems such as flaviviruses, employ in order to deliver their genomes to initiate infection. With structural elucidation of these steps, it becomes possible to identify key stages for targeting by inhibitors with means to understand their mechanisms. For class-I fusion systems such as HIV, neutralizing antibodies have been described that bind to intermediate forms of its fusion proteins and can potentially arrest the fusion process ^50; 51^. Few examples of antibodies that trap flaviviruses in an intermediate stage that prevents fusion have also been identified ^52; 53^. With better understanding of the key structural stages in alphavirus membrane fusion, as probed in this work, it may be possible to develop better strategies to inhibit theses viruses’ fusion and entry. At a broader level, resolving the molecular processes of CHIKV fusion also advances our understanding of fundamental aspects in protein-mediated membrane fusion, which is an essential biological process involved not only in enveloped virus infection but also in cell-to-cell fusion, intracellular vesicle fusion, gamete fusion and synaptic vesicle signaling.

## Methods

### CHIKV preparation and purification

CHIKV strain S27 was propagated and purified similar to previous reports ^33^. Briefly, BHK-21 (Baby Hamster Kidney) cells were cultured at 37C and 5% CO2 in Dubelcco’s minimal essential medium (DMEM) supplemented with 10% FBS (fetal bovine serum). Cells were infected with virus at m.o.i of 4.0 and virus particles allowed to infect for 1.5 hours. After 25-27 hours post-infection, the medium was collected, and virus particles were pelleted by ultracentrifugation for 2 hours at 19000rpm in a Beckman Type 19 rotor at 4ºC. The pelleted virus was resuspended overnight in HNE buffer (50mM HEPES, 150mM NaCl and 0.1mM EDTA), pH 7.4. The resuspended sample was applied to a sucrose gradient. The sucrose gradient was spun in an ultracentrifuge at 20,000rpm in a Beckman SW41 rotor overnight at 4°C. The virus band was extracted, inactivated by exposure to UV lamp, aliquoted and snap frozen in liquid nitrogen. Prior to experiments, inactivated CHIKV in 40% sucrose solution was dialyzed into HBS (10mM HEPES, 150mM NaCl, 50mM sodium citrate) buffer pH7.5, for 4-6 hours at 4ºC.

### Liposome preparation

Liposomes composed of phosphatidyl choline (PC), phosphatidyl ethanolamine (PE), sphingomyelin and cholesterol (molar ratio 1:1:1:1.5) were prepared by lipid extrusion method described previously^30; 33^. Stock solutions of the different components were prepared in chloroform and combined in appropriate ratios. The combined lipid solutions were dried under nitrogen gas. The lipid films were then resuspended in HBS (10mM HEPES, 150mM NaCl, 50mM sodium citrate (pH 7.5) and passed through five liquid nitrogen freeze-thaw cycles. For thaw cycles, water bath at 50ºC was used. The resuspended solution was extruded 21 times through a 200-nm polycarbonate membrane. All lipids and membrane were purchased from Avanti Polar Lipids. The resulting liposomes were passed over a PD-10 desalting column (GE Healthcare) and stored in pH 6.0 HBS buffer.

### Sample preparation and data collection for single particle cryo-EM

Inactivated CHIKV in HBS buffer (pH 7.5) was applied to lacey carbon grids with a thin continuous carbon film (400 mesh) (Electron Microscopy Sciences). The grids were glow discharged (negative charge) under vacuum using 25mA current for 30 seconds. A 3µl aliquot of the sample was applied to these grids at 4ºC and 100% humidity, blotted for 3seconds and immediately plunge frozen in liquid ethane using a Vitrobot Mark IV (FEI Co.).

Vitrified grids were imaged using a 300kV Titan Krios (FEI Co.) equipped with a K2 Summit direct electron detector (Gatan Inc.) and post-specimen energy filter. Micrographs were collected at a nominal magnification of 105000X with a corresponding pixel size of 1.35 Å/pixel in counting mode. A dose rate of ∼8 e^−^/pixel/s was used with 200 ms exposure per frame and 50 frames per image. Data was collected with a defocus range from 1.5 to 3.5 µm. A total of 495 micrographs were collected using the automated data collection software Leginon ^54^.

### Single particle cryo-EM data processing and structure determination

All data processing steps were carried out within the Relion software package^55; 56^. Frame alignment and dose-weighting was done using MotionCor2^57^. CTF estimation was performed using CTFFIND4^58^. A total of 7741 particles were picked automatically using 2D reference templates. Particles were extracted at 4x binning and the binned particle stack was used for unsupervised 2D classification. Further processing with 3D classification did not produce any individually better class. A total of 5806 selected particles were thus used for 3D refinement with icosahedral symmetry imposed. Two initial models, a sphere and a low pass filtered reconstruction of CHIKV virus-like particle (EMD-5580), were used for separate 3D refinement runs. Both refinements converged to near identical structures of CHIKV. The particle stacks were unbinned progressively for further refinements. Map sharpening and post-processing was also carried out in Relion which gave a final structure with resolution 6.75 Å using the “gold-standard” FSC cutoff of 0.143.

### Sample preparation for cryo-ET

To make grids for cryo-ET, 400 mesh Lacey carbon grids with a layer of ultrathin carbon (Electron Microscopy Sciences) were glow discharged for 30 seconds. A 3µl aliquot of CHIKV sample was mixed with 10nm gold beads (Aurion BSA Gold Tracer 10nm) at a ratio of 15:1 (v/v). The sample was allowed to adsorb on the grid at room temperature for 15 seconds. Pre-calculated volumes of liposome mixture along with appropriate volume of HBS pH 3.0 were then added to the grids to drop the pH to desired values. Grids were then incubated at room temperatures for different time points inside a humidity-controlled chamber to avoid evaporation. Sample grids were then loaded onto a Vitrobot Mark IV (FEI Co.) at 4ºC and 100% humidity, blotted for 7-8seconds and plunge frozen in liquid ethane.

### Cryo-ET Data collection

Frozen grids were imaged using a 300 kV Titan Krios with a Gatan K2 Summit direct electron detector and GIF energy filter with slit width of 20 eV. Tilt-series were collected in a dose-symmetric tilting scheme from −60° to +60° or from −54° to +54° with a step size of 3° using Leginon ^54^ or SerialEM softwares ^59^. Tilt-series were collected either in counting mode at a magnification of 81000X, corresponding to a pixel size of 1.69 Å per pixel or in super-resolution mode at a magnification of 53000X, corresponding to a pixel size of 0.8265 Å per pixel. The total dose per tilt series ranged between ∼60-80 e^-^/Å^2^. A total of 441 tilt-series were collected across multiple sessions.

### Tomogram reconstruction

Tilt-series image frames were corrected for electron beam-induced motion using Motioncor2^57^. Tilt images were then processed using batch tomography in IMOD^60^ using standard procedures to generate 3-dimensional tomogram reconstructions. Tilt-series images were aligned using the gold bead markers. The aligned images were then reconstructed to give a 3D volume using weighted back-projection. The final tomograms were binned, low pass-filtered and contrast enhanced in ImageJ for visualization^61^. Supplemental tomogram movies were also made using ImageJ with pixel size in direction perpedicular to electon beam (x-y direction) being 10.14 Å/pixel and 50.7 Å/pixel in the direction of the electron beam (z-direction). Volumes were rendered in 3D using UCSF Chimera^62^.

### Sub-tomogram averaging

#### Low pH CHIKV virions and post-fusion nucleocapsids

Tilt-series from pH 5.9, 5.1 and 5.1 at 30 seconds to 1 minute timepoints were imported into EMAN2’s sub-tomogram averaging pipeline ^63^. 1k X 1k tomograms were generated within EMAN2 using default parameters. A total of 70 unattached or mildly attached virus particles were picked manually in the e2spt_boxer.py interface ^63^. Sub-volumes were extracted at 8xbinning corresponding to a pixel size of 6.612 Å per pixel. Sub-tomogram alignment and refinement was carried using a spherical mask that covers an entire virus particle. A ring mask that encompasses only the outer glycroprotein shell and membrane was also tried. Different initial models, low pass filtered sphere map or CHIKV map or initial model generated within EMAN2 using stochastic gradient descent principle, were used as separate starting points. In all the cases, the output map did not converge to any with interpretable density features. A smaller radius mask covering only the nucleocapsid region was also attempted to check if the nucleocapsid in low pH CHIKV virions retained its neutral-pH structure. These attempts also failed to give any interpretable density map structure. Using EMAN2 tools, 2D radial density average plot of the sub-tomogram averaged CHIKV virion map was calculated for analysis.

A similar protocol as above was used for calculating the post-fusion nucleocapsid structure. Tilt-series from later fusion timepoints were imported into the EMAN2 pipeline. Nucleocapsids released into the liposome lumen were manually picked in the e2spt_boxer.py interface ^63^. A total of 122 sub-volumes that appeared reasonably spherical were extracted at 8xbinning corresponding to a pixel size of 6.612 Å per pixel. A spherical mask covering the entire nucleocapsid particle was used. Different initial models, low pass filtered sphere map or CHIKV nucleocapsid structure or initial model generated within EMAN2 using stochastic gradient descent principle, were used as separate starting points. In all cases, the output map did not converge to any interpretable density features.

### Post-fusion E1 glycoprotein

A total of 40 tomograms from late time-points that contained fused virions with distinguishable protein features on the external surface of liposomes were selected. Ctf-estimation for the tilt-series was carried out in EMAN2^63^ and ctf-correction applied using ctfplotter in IMOD^64^. Protein spikes on surface of liposomes were picked manually in 3d-mod^65^. Each protein unit was identified using two points, with first point placed distal to the membrane and the second point placed at the protein end close to the membrane. Using these points, motive lists with coordinate positions and rotation angles with respect to the designated ‘y’ axis of the tomogram was calculated for each particle using the ‘stalkInit’ program within the PEET software suite^66^. A total of 591 protein spikes were picked. Subsequent sub-tomogram volume extraction, alignment and averaging was also carried out within PEET using binned data corresponding to pixel size of 6.612 Å per pixel. A soft cylindrical mask that contained the protein spike and outer membrane layer was used. Search parameters allowed for a complete 360º search along the long axis of the protein spikes but restricted the search in the other two directions to ±60° with 0° being the long axis of the protein spikes. Initial coarse searches were followed by progressively finer search parameters. Duplicate removal was enabled to weed out overlapping volumes. Two initial models were tested -- a randomly selected sub-volume and a generated map of the post-fusion E1 glycoprotein trimer structure from Semliki Forest virus (low pass filtered to 60Å). Missing wedge compensation was also applied within PEET during the alignment and averaging process. After initial alignment and averaging using standard averaging parameters as suggested by the PEET tutorials, the output sub-tomogram averages from the jobs with different initial models had similar structures. No symmetry was imposed in the initial steps. In the output sub-tomogram average of the spike using all particles, 3-fold symmetry was observed along the long axis of the spike. The dataset was spilt into even and odd datasets and averaged separately. Three-fold symmetry was applied during the averaging routine to give a final density map of resolution of 27.2 Å at 0.5 FSC cut-off. Crystal structure of the Semliki Forest virus post-fusion E1 glycoprotein trimer (PDB: 1RER) was fitted into the map density using UCSF Chimera.

## Acknowledgements

We thank the University of Washington Arnold and Mabel Beckman Cryo-EM for data collection time and support. A portion of this research was supported by NIH grant U24GM129547 and performed at the PNCC at Oregon Health and Science University (OHSU), a DOE Office of Science User Facility sponsored by the Office of Biological and Environmental Research, U.S.A. We also thank Dr. Mareike K. S. van Duijl-Richter, University of Groningen, Netherlands, for her help in virus production. This work was supported by NIH grant R01-GM099989 (KKL).

## Author Contributions

V.M.P.: Conceptualization, Formal analysis, Investigation, Validation and Writing.: J.S.B.: Conceptualization, Resources.: J.M.S.: Resources, Writing review. K.K.L.: Conceptualization, Resources, Writing, Funding acquisition.

## Data Availability

Sub-tomogram averaged density map of post-fusion E1 trimer with corresponding fitted atomic model has been deposited with accession codes EMD-XXXX and PDB ID XXXX.

