## Supplemental Information for "Visualization of conformational changes and membrane remodeling leading to genome delivery by viral class-II fusion machinery"

Affiliations:

**Supplementary Figures:**

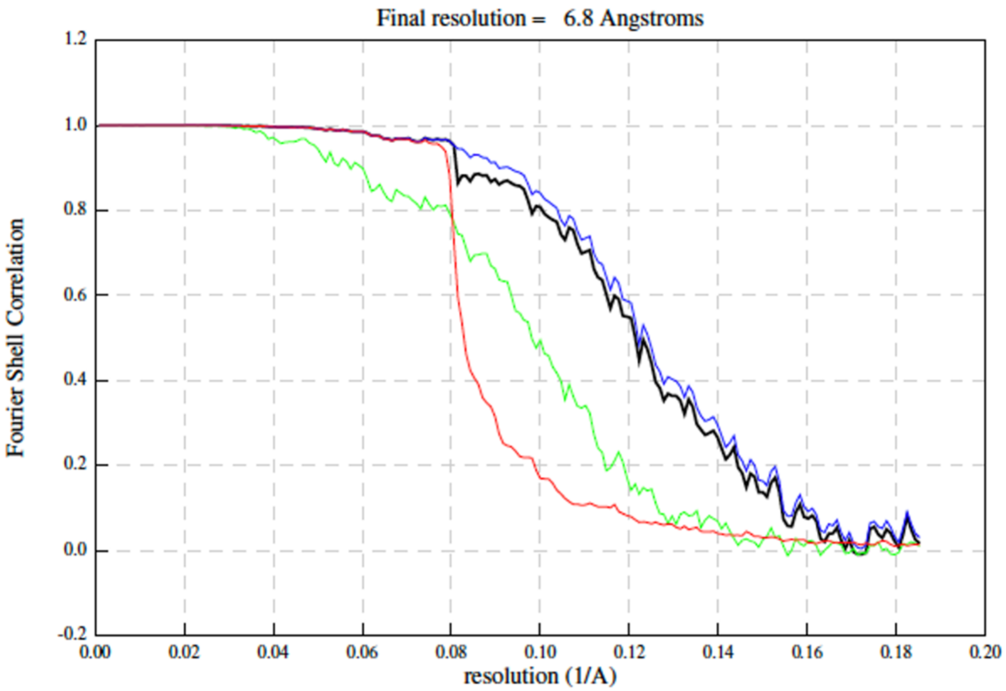

**Supplementary Figure 1:** Gold-standard Fourier shell correlation (FSC) curves. Lines represent correlation corrected (black), masked (blue), unmasked (green), and phase randomized (red) Fourier Shell correlation curve maps. Resolution at "gold-standard" FSC cutoff of 0.143 is given as final resolution.

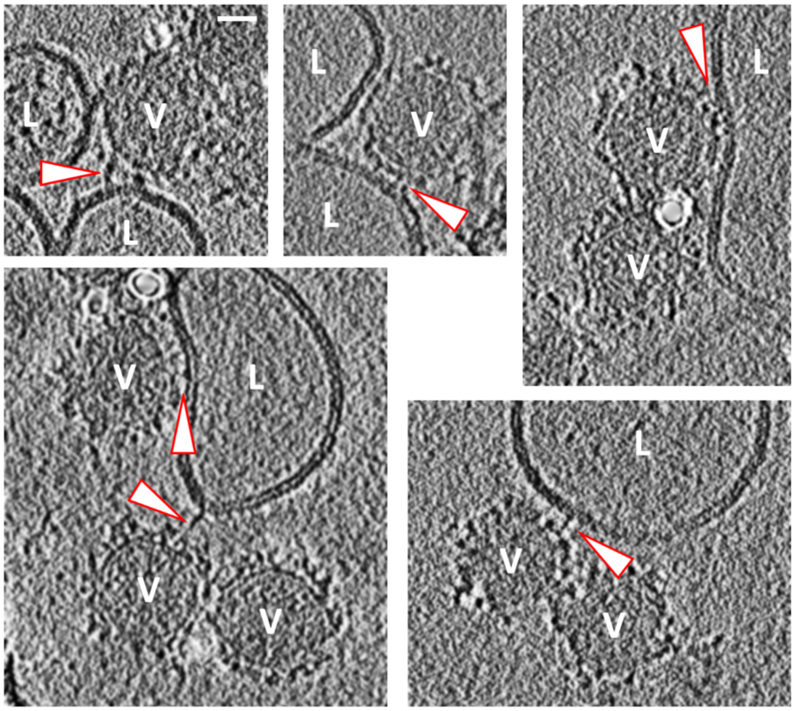

**Supplementary Figure 2:** CHIKV-liposome interactions at pH >6.0. Discrete densities (arrow heads) connecting CHIKV (V) to surrounding liposomes (L) is seen at pH 6.3 (top panels) and pH 6.1 (bottom panels). Black is high density. Scale bar is 200 Å in length.

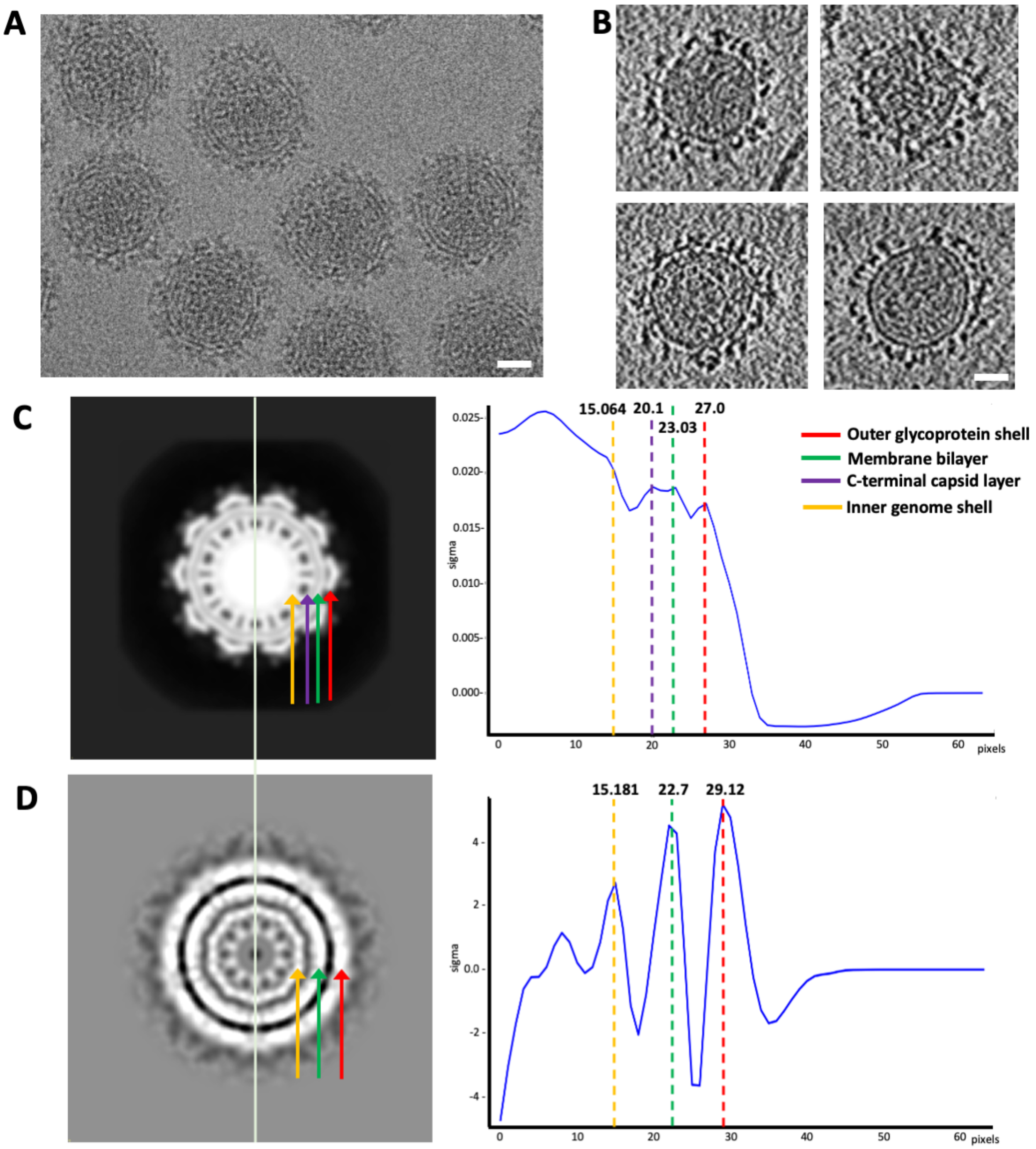

**Supplementary Figure 3:** Sub-tomogram averaging analysis of low pH (<6.0) treated CHIKV. A. EM micrograph of neutral pH CHIKV. B. Tomogram cross-section of low pH treated CHIKV. In panels A and B, black is high density and scale bar is 200 Å. C. Left: Cross-section of neutral pH CHIKV EM density map low pass filtered to 45Å resolution. Right: 2D radial density plot of the same. D. Left: Cross-section of low pH treated CHIKV sub-tomogram averaged EM density map at 45Å resolution showing lack of discernable protein features (as compared to panel C). Right: 2D radial density plot of the same. In both panels C and D, red arrow or dotted line indicates the outer glycoprotein shell, green arrow or dotted line indicates the membrane bilayer, purple arrow or dotted line indicates the nucleocapsid shell juxtaposed underneath the viral membrane, yellow arrow or dotted line indicates the outer radius of the central genome region. The 2D radial density plot shows that though the internal genome region and membrane bilayer are at similar radii, the glycoprotein shell peak is at a marginally higher radius in the low pH treated virion when compared to the neutral pH CHIKV. Pixel size of density maps and related plots are 10.14 Å/pixel.

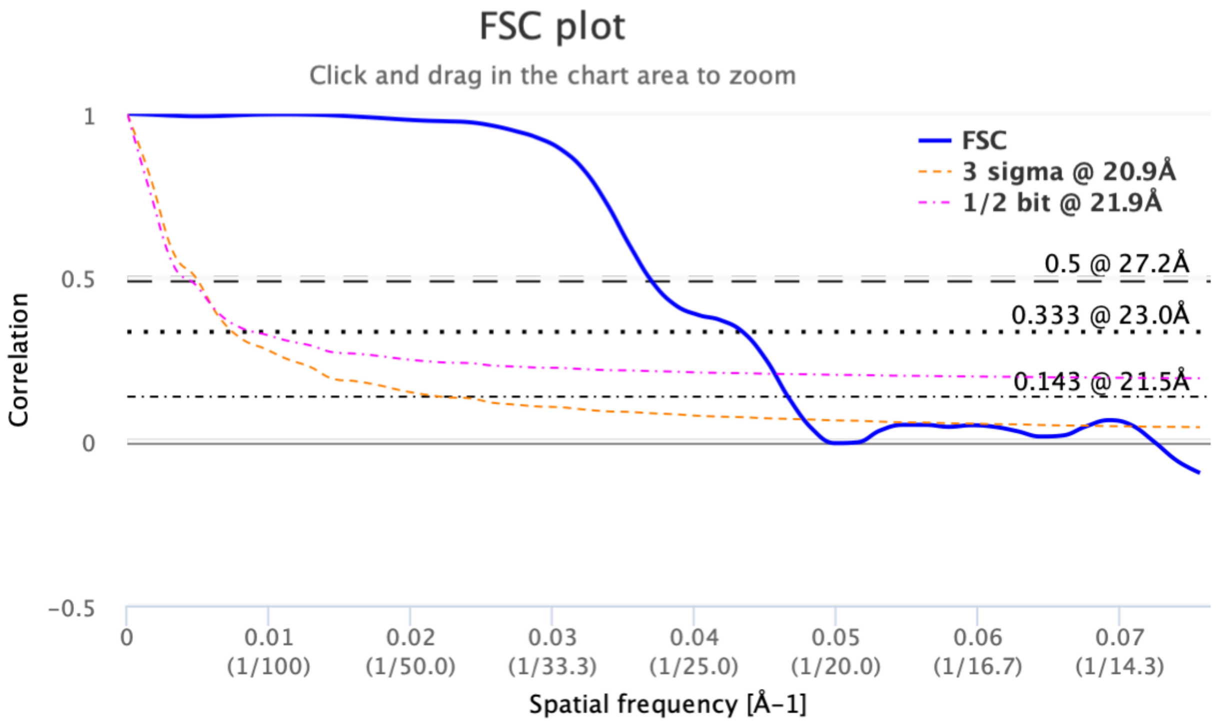

**Supplementary Figure 4:** Fourier Shell correlation (FSC) curve. FSC curve computed using even and odd half-maps calculated for the sub-tomogram averaged post-fusion E1 trimers. Plot was calculated using the Electron Microscopy DataBank (EMDB) FSC server.

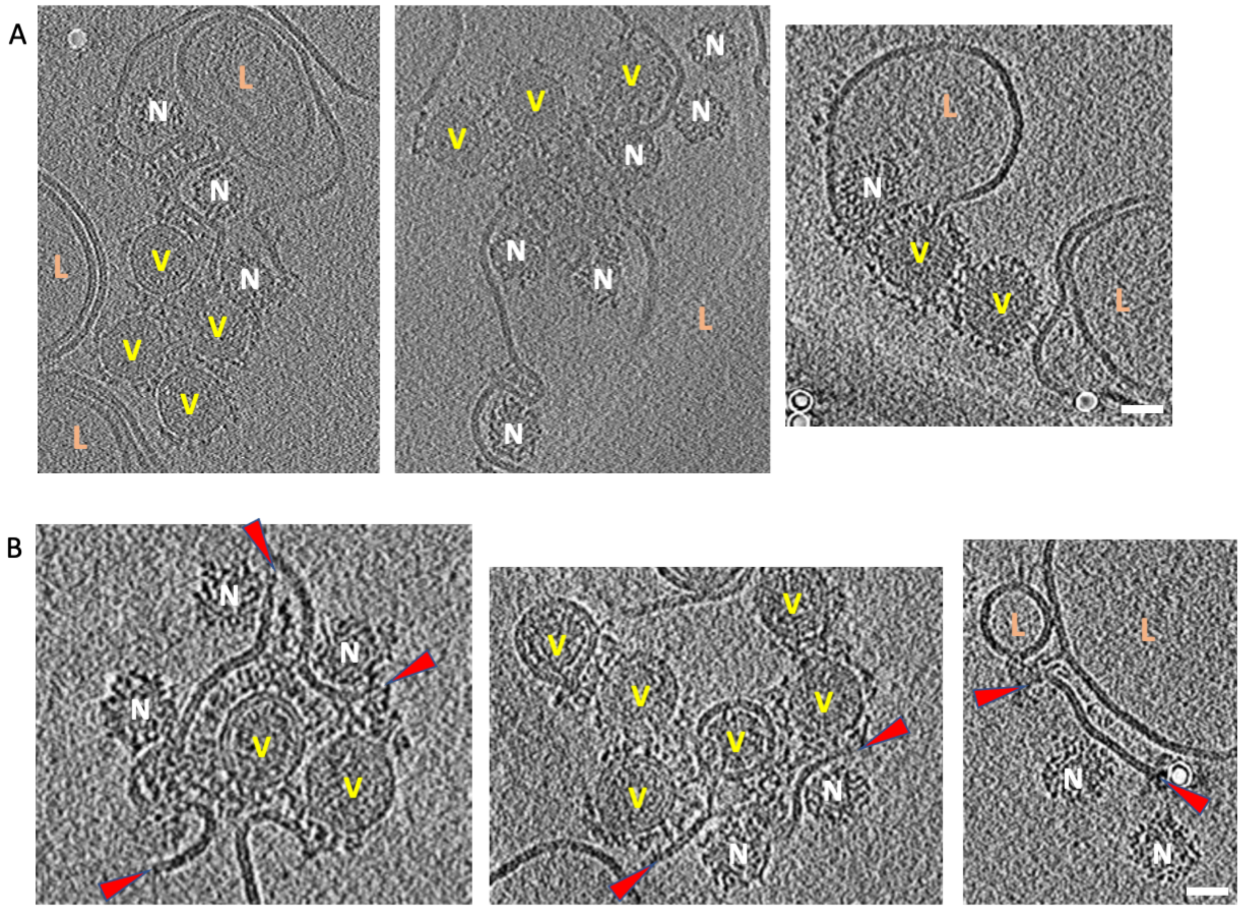

**Supplementary Figure 5:** Effect of prolonged exposure at low pH (pH 5.6 and pH 5.1) on CHIKV. A. Tomogram cross-sections showing examples of multiple CHIKV aggregates fusing with liposomes. B. Tomogram cross-sections showing examples of CHIKV fusing with other virions and releasing its nucleocapsids. Red arrowheads indicate the ends of the fused virus membranes. In all panels: Black is high density. Virus is denoted as V, liposomes as L and nucleocapsids as N. Scale bars are 300Å in length.

Supplementary movies legends:

**Supplementary movie 1**: **Stage I-membrane recruitment**. Cryo-electron tomography reconstruction of CHIKV-liposome interaction showing an example of stage I – membrane recruitment. CHIKV is indicated as V, liposome as L and interaction site is indicated by a yellow arrow. Scale bar is 200 Å. Black is high density. For all supplemental movies, tomogram pixel size in direction perpedicular to electon beam (x-y direction) is 10.14 Å/pixel and 50.7 Å/pixel in the direction of the electron beam (z-direction).

**Supplementary movie 2**: **Stage II-membrane attachment.** Cryo-electron tomography reconstruction of CHIKV-liposome interaction showing an example of stage II – membrane attachment. CHIKV is indicated as V, liposome as L and interaction site is indicated by yellow arrows. Green arrow indicates the presence of a stage III – E1-HT example in the same virion. Scale bar is 200 Å. Black is high density.

**Supplementary movie 3**: **Stage III-E1-HT formation.** Cryo-electron tomography reconstruction of CHIKV-liposome interaction showing an example of stage III – formation of E1-HT. CHIKV is indicated as V, liposome as L and interaction site is indicated by yellow arrow. Green arrow indicates the presence of a stage IV – E1-HT membrane insertion example in an adjacent virion. Scale bar is 200 Å. Black is high density.

**Supplementary movie 4**: **Stage IV-E1-HT membrane insertion.** Cryo-electron tomography reconstruction of CHIKV-liposome interaction showing an example of stage IV –E1-HT membrane insertion. CHIKV is indicated as V, liposome as L and interaction site is indicated by yellow arrow. Scale bar is 200 Å. Black is high density.

**Supplementary movie 5**: **Stage V-opposing membrane superposition.** Cryo-electron tomography reconstruction of CHIKV-liposome interaction showing an example of stage V – opposing membrane superposition. CHIKV is indicated as V, liposome as L and interaction site is indicated by yellow arrow. Scale bar is 200 Å. Black is high density.

**Supplementary movie 6**: **Stage VI-tight membrane apposition.** Cryo-electron tomography reconstruction of CHIKV-liposome interaction showing an example of stage VI – tight membrane apposition. CHIKV is indicated as V, liposome as L and interaction site is indicated by yellow arrow. Scale bar is 200 Å. Black is high density.

**Supplementary movie 7**: **Stage VII-hemifusion.** Cryo-electron tomography reconstruction of CHIKV-liposome interaction showing an example of stage VII – hemifusion. CHIKV is indicated as V and liposome as L. Hemifusion site is indicated by yellow arrows. Scale bar is 200 Å. Black is high density.

**Supplementary movie 8**: **Stage VIII-fusion pore formation.** Cryo-electron tomography reconstruction of CHIKV-liposome interaction showing an example of stage VIII – fusion pore formation. CHIKV is indicated as V, liposome as L and interaction site is indicated by yellow arrow. Scale bar is 200 Å. Black is high density.

**Supplementary movie 9**: **Stage IX-nucleocapsid release.** Cryo-electron tomography reconstruction of CHIKV-liposome interaction showing an example of stage IX – nucleocapsid release. Liposome is labeled as L. Yellow arrows indicate two post-fusion nucleocapsids released into the liposome lumen. Green arrows denote post-fusion E1 glycoprotein trimers distributed on the liposome membrane. Scale bar is 200 Å. Black is high density.

Supplemental Table 1: Number of CHIKV-liposome contacts identified for each stage in the membrane fusion process for different pH and timepoints.

| | | E1-attachment | | E1 HT/Membrane Insertion | | | Hemifusion | | Post-fusion NC-release | |
|---|---|---|---|---|---|---|---|---|---|---|
| **pH** | **Time (minutes)** | **Recruitment** | **Attachment** | **E1-HT** | **E1-HT insertion** | **Membrane superposition** | **Hemifusion** | **Fusion pore** | **Post-fusion** | **Total** |
| 5.1 | 0.5 | 0 | 16 | 8 | 2 | 6 | 0 | 1 | 1 | 34 |
| | 1 | 5 | 17 | 5 | 8 | 2 | 2 | 5 | 19 | 63 |
| | 3 | 0 | 48 | 12 | 11 | 12 | 9 | 10 | 64 | 166 |
| 5.6 | 0.5 | 100 | 50 | 2 | 0 | | 1 | 0 | 0 | 153 |
| | 2 | 3 | 43 | 10 | 17 | 2 | 0 | 2 | 2 | 79 |
| | 4 | 0 | 6 | 3 | 5 | 4 | 1 | 3 | 27 | 49 |
